# Screening of Cellular Senescence (CS) Related Genes as Biomarkers and Therapeutic Targets for Glioblastoma (GBM) by Integrated Machine Learning (IML)

**DOI:** 10.1101/2025.08.19.670890

**Authors:** Tianle Xue, Yufan Hu, Haoju Lyu, Tian Xie

## Abstract

Glioblastoma (GBM) is an aggressive brain tumor with limited prognostic biomarkers and therapeutic targets. This study applied an integrated machine learning (IML) framework to discover cellular senescence (CS)-related gene biomarkers and candidate therapeutic targets in GBM. Using Gene Expression Omnibus (GEO) training and validation cohorts, 113 machine learning models across 11 algorithms were integrated to pinpoint CS-associated gene signatures. Differential expression analysis combined with overlap of a CellAge senescence gene set yielded 129 CS-related differentially expressed genes (DEGs). Functional enrichment of these DEGs highlighted pathways related to cellular senescence and cell cycle regulation (Gene Ontology and KEGG). A multivariate classifier constructed via stepwise generalized linear modeling (GLM) and LASSO achieved high diagnostic performance (area under the ROC curve 0.92 in training, each independent validation set is over 0.85). Seven top-ranked genes from this model were validated, with TGFβI emerging as the most robust biomarker (AUC > 0.85) and its elevated expression was associated with significantly shorter overall survival. Spatial transcriptomics and single-cell RNA sequencing localized TGFβI expression to tumor cell clusters harboring high copy number variation (CNV) burdens. Immune microenvironment profiling linked TGFβI expression with increased macrophage infiltration. Finally, single-cell gene set enrichment (scGSEA) and AUCell analyses indicated enrichment of ECM–receptor interaction signaling in TGFβI-expressing cells. In summary, IML combined with spatial and single-cell transcriptomics identified TGFβI as a potent CS-related biomarker and a promising therapeutic target in GBM.

## Introduction

Glioblastoma (GBM) is the most prevalent and aggressive primary malignant brain tumor in adults, accounting for approximately 49% of such cases^[1]^. Its incidence has been on the rise, with a notable increase observed in recent years, and predominantly affects individuals aged 45 to 70, with a higher occurrence in males compared to females ^[2, 3]^. The etiology of GBM remains largely unclear, however, certain risk factors have been identified. Exposure to ionizing radiation and genetic predispositions, such as Li-Fraumeni syndrome and neurofibromatosis, have been implicated in its development^[4]^. Despite advances in multimodal treatment approaches, comprising surgical resection, radiotherapy, and chemotherapy, the prognosis for GBM remains poor^[5]^. The median survival time is approximately 12 to 15 months, with a five-year survival rate below 10%^[6]^. The aggressive nature of GBM, coupled with its resistance to conventional therapies and high recurrence rate, underscores the urgent need for novel diagnostic and therapeutic strategies. Identifying reliable biomarkers and therapeutic targets is crucial for improving patient outcomes.

Cellular senescence is a state of irreversible cell cycle arrest triggered by various stressors, including DNA damage, oncogene activation, and telomere attrition^[7]^. Senescent cells exhibit distinct phenotypic changes, such as enlarged morphology, altered gene expression, and the secretion of pro-inflammatory cytokines, collectively known as the senescence-associated secretory phenotype (SASP). While senescence acts as a tumor suppressive mechanism by preventing the proliferation of damaged cells, accumulating evidence suggests that the SASP can promote tumor progression and metastasis by altering the tumor microenvironment^[8]^. Glioblastoma (GBM), the most aggressive primary brain tumor, often exhibits features of cellular senescence. Studies have shown that GBM cells can enter a senescent-like state in response to therapy, contributing to treatment resistance and tumor recurrence^[9]^. However, the complex role of senescence in GBM pathogenesis remains poorly understood, necessitating comprehensive analyses to identify senescence-related biomarkers and therapeutic targets.

To address this gap, we utilized CellAge, a curated database of human genes associated with cellular senescence^[10]^. By integrating machine learning approaches, we screened for senescence-related genes that may serve as potential biomarkers and therapeutic targets for GBM, aiming to enhance our understanding and treatment of this devastating disease.

Machine learning (ML) has revolutionized cancer diagnostics by enabling the analysis of high-dimensional genomic data to identify disease-specific gene expression patterns. Studies have demonstrated ML’s efficacy in classifying various cancers based on gene expression profiles^[11]^, achieving high accuracy rates. Beyond diagnosis, ML techniques facilitate the discovery of therapeutic targets by sifting through complex oncogenomic datasets to pinpoint key driver genes and pathways^[12]^. Integrated machine learning (IML), which combines multiple algorithms and multi-modal biomedical data (e.g., multi-omics and clinical features), offers even greater advantages. IML approaches leverage diverse data sources – from genomics to pathology – yielding more robust and generalizable biomarkers and targetable vulnerabilities than single-modality analyses^[13]^. Harnessing IML in GBM is therefore poised to uncover critical cellular senescence–associated gene signatures implicated in tumor progression^[14]^, potentially revealing new biomarkers and therapeutic targets for this lethal cancer.

In this study, we integrated 11 different machine learning algorithms, generating 113 machine learning models through their individual models or combinations. By using these 113 machine learning models to screen potential targets about the cellular senescence related genes, the accuracy of the results will be significantly improved. At the same time, it can make the disease-related genes obtained by analysis more reliable. The whole working flow is demonstrated in Figure 1.

**Figure 1.**
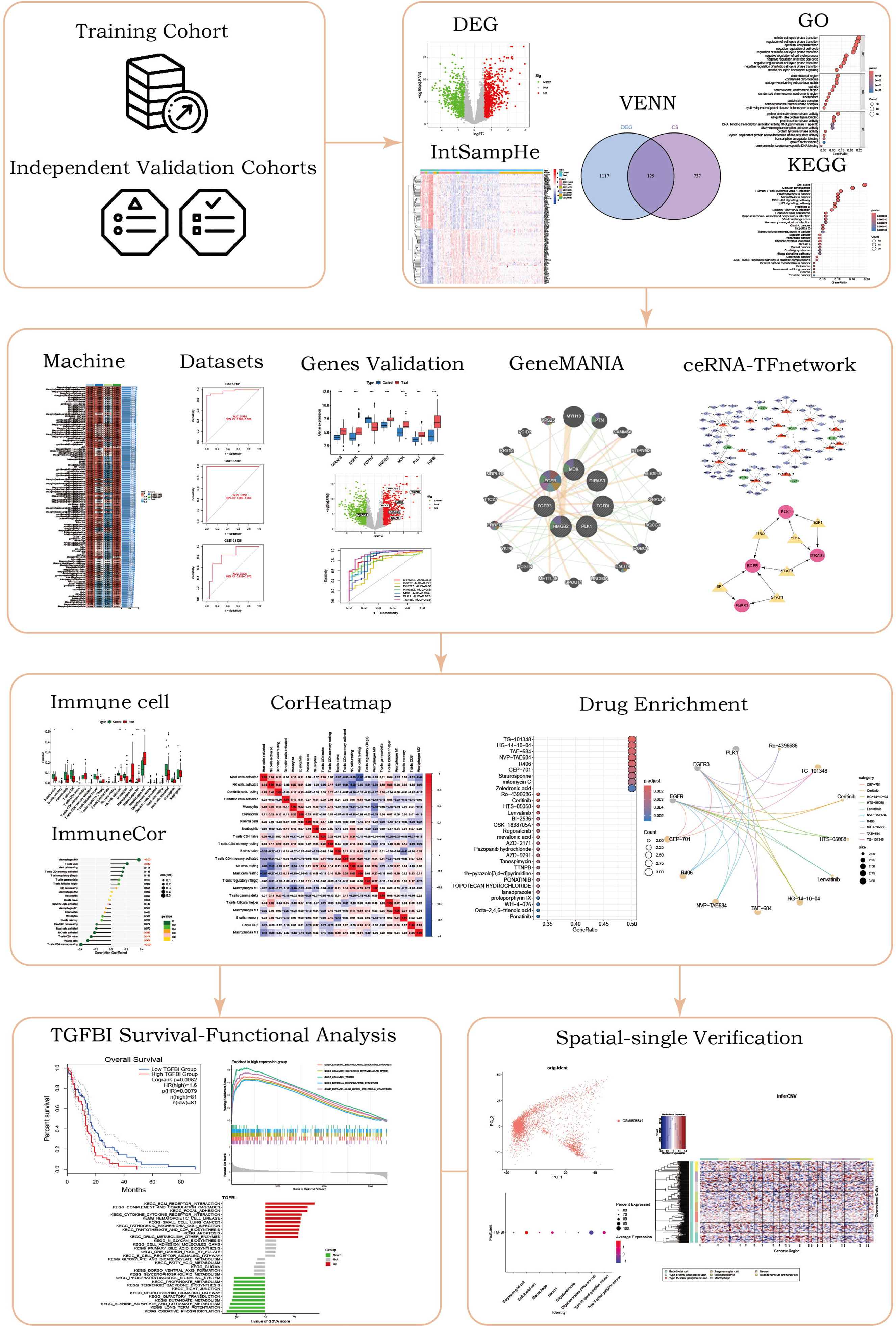
Workflow for identifying Cellular Senescence (CS)-related biomarkers and therapeutic targets in Glioblastoma (GBM). Gene expression data from training and independent cohorts underwent differential expression analysis (DEG), integration, and intersection to pinpoint CS-related genes. Functional enrichment (GO and KEGG) characterized their biological roles. Integrated machine learning validated key gene signatures, with network analyses (GeneMANIA, ceRNA, transcription factors) revealing gene interactions. Immune profiling, drug enrichment, survival analyses, and functional characterization (GSEA, GSVA) highlighted TGFβI as a robust biomarker linked to poor survival and ECM receptor interactions. Spatial transcriptomics and single-cell analyses further validated TGFβI within malignant clusters with elevated chromosomal copy number variations (CNV), emphasizing its potential as a therapeutic target.

## Methods

### 1. Data Acquisition and Preprocessing for Screening CS-Related Genes in GBM

Seven publicly available microarray gene expression datasets (GSE122498, GSE12657, GSE13276, GSE15824, GSE22866, GSE22867, GSE30563) were obtained from the NCBI Gene Expression Omnibus (GEO) database and combined as a training cohort, while three additional GEO datasets (GSE50161, GSE137901, GSE161528) served as an independent validation cohort. All data were processed using probe identifiers were converted to gene symbols based on platform annotation files. The combined expression data were transformed if necessary and then normalized across arrays using the limma package to ensure comparable expression distributions. Finally, batch effects among the merged datasets were corrected using the ComBat method implemented in the sva package for subsequent integrated machine learning analyses of cellular senescence (CS)-related genes in GBM.

### 2. Assessment of Batch Effect Correction

All the training set for this study were merged and batch-corrected to minimize cross-dataset technical variability. Batch effect removal was then evaluated using boxplots and principal component analysis (PCA). Boxplots of gene expression values across samples (grouped by dataset) before and after correction revealed more consistent distributions post-correction. Similarly, PCA was performed to visualize clustering among samples from different datasets; after correction, samples no longer grouped by their dataset of origin, indicating that batch-associated variance was effectively reduced.

### 3. Identification of Differentially Expressed Cellular Senescence-Related Genes

Gene expression profiles from seven training datasets comprising GBM and normal brain samples were integrated for analysis. Differentially expressed genes (DEGs) between GBM and normal samples were identified using the limma R package^[15]^, considering as significant those with |log2 fold change (FC)| > 0.585 and adjusted p-value < 0.05. The resulting DEGs were then intersected with known cellular senescence-associated genes from the CellAge database to yield a set of candidate CS-related DEGs for further analysis^[16]^.

### 4. Functional Enrichment Analysis

Functional enrichment analysis of the identified cellular senescence (CS)-related differentially expressed genes (DEGs) was performed using standard bioinformatics tools, notably the clusterProfiler R package^[17]^. Gene Ontology (GO) term and Kyoto Encyclopedia of Genes and Genomes (KEGG) pathway enrichment analyses were carried out. GO enrichment covered the Biological Process (BP), Cellular Component (CC), and Molecular Function (MF) categories. Significance thresholds of p-value < 0.05 and adjusted p-value < 1 were applied to identify significantly enriched GO terms and pathways.

### 5. Integrated Machine Learning Based Gene Selection

We applied an integrated machine learning framework to identify cellular senescence (CS)-related genes as potential biomarkers and therapeutic targets in glioblastoma (GBM). A total of 113 predictive models were constructed by applying 12 different algorithms individually and in all pairwise combinations. The algorithms evaluated included Lasso, Ridge, stepwise GLM, Random Forest, XGBoost, Elastic Net, LDA, plsRglm, generalized boosted regression modeling, Naive Bayes, GLMBoost, and SVM. Each model’s predictive performance was evaluated using the concordance index (C-index), and the model achieving the highest C-index was selected as optimal. Genes selected by this top-performing model were designated as candidate CS-related biomarkers and therapeutic targets for GBM. All model development and evaluation steps were conducted using standard machine learning software libraries to ensure reproducibility of results.

### 6. Validation of Diagnostic Efficacy and Differential Expression of Candidate Genes

Receiver operating characteristic (ROC) curve analysis was performed to validate the diagnostic performance of the candidate CS-related gene signature. ROC curves were constructed for both training and validation cohorts, and the area under the curve (AUC) was measured to quantify predictive accuracy. Differential expression analysis between GBM and normal brain samples identified significantly dysregulated genes, and the candidate genes from the top-performing model were highlighted on a volcano plot. Expression levels of these candidates in tumor versus normal tissues were visualized with boxplots, and group differences were assessed using the Wilcoxon rank-sum test. Individual ROC curves were also plotted for several top-ranked genes to evaluate single-gene diagnostic utility.

### 7. Gene Interaction Network Analysis

All cellular senescence (CS)-related candidate genes identified by the optimal machine learning model were submitted to GeneMANIA to construct a gene–gene interaction network^[18]^. This network was used to visualize functional associations between the candidate genes and their interacting partners.

### 8. Construction of ceRNA and Transcription Factor Regulatory Networks

Based on the genes identified via integrated machine learning as potential biomarkers and therapeutic targets for glioblastoma (GBM), competing endogenous RNA (ceRNA) and transcription factor (TF) regulatory networks were constructed. For the ceRNA network, candidate mRNAs were linked to corresponding long non-coding RNAs (lncRNAs) and microRNAs (miRNAs) based on interactions predicted using starBase v2.0 (https://starbase.sysu.edu.cn/) with thresholds of prediction score >0.8 and p-value <0.05. Additionally, transcription factors targeting candidate genes were identified through the TRRUST database (https://www.grnpedia.org/trrust/), retaining interactions with confidence scores >0.7. The resulting ceRNA and TF-gene interactions were visualized and analyzed using Cytoscape v2.0 software (Shannon et al., 2003, https://cytoscape.org/). Network parameters such as node degree and betweenness centrality were computed to identify key regulatory elements. These regulatory networks were constructed to clarify potential molecular mechanisms underlying the roles of cellular senescence-related genes in GBM.

### 9. Immune Cell Profiling and Correlation With Candidate Genes

We applied the CIBERSORT algorithm to normalized gene expression profiles of GBM tumor and normal brain samples to estimate the relative abundances of 22 infiltrating immune cell types^[19]^. Differences in immune cell composition between GBM and normal groups were visualized using bar plots and boxplots. We then performed correlation analysis to examine associations between the expression levels of the most relevant CS-related candidate genes and the corresponding immune cell fractions. Significant gene-immune correlations were illustrated using lollipop plots and network diagrams to highlight the relationships between candidate genes and immune cell types. These analyses were conducted to explore the potential immunological relevance of the identified candidate genes in GBM.

### 10. Drug Enrichment Analysis of Candidate Targets

Drug enrichment analysis was performed using the clusterProfiler R package^[17]^ to identify potential therapeutic compounds targeting the candidate CS-related genes. The top-ranked CS-related genes from the integrated machine learning (IML) analysis served as input, with drug–gene associations obtained from the Drug Signatures Database (DSigDB). Enriched compounds were identified using a hypergeometric test, with significance defined at p < 0.05 and Benjamini–Hochberg adjusted p < 0.05.

### 11. Survival Analysis of Candidate Genes Using GEPIA2

Overall survival (OS) analyses of candidate genes were performed using the GEPIA2 web server (http://gepia2.cancer-pku.cn/), which utilizes TCGA GBM patient data. GEPIA2 automatically stratifies patients into high-and low-expression groups based on each gene’s median expression level and generates Kaplan–Meier survival plots with hazard ratios and log-rank P-values. The gene displaying the most relevance in OS between high and low expression groups (lowest log-rank P value) was identified for further analysis. All Kaplan–Meier survival plots and associated statistics were generated automatically by the GEPIA2 platform^[20]^.

### 12. Functional Enrichment Analysis Based on Prognostic Gene Expression

The top prognostic gene identified from survival analysis was selected for downstream functional enrichment analysis. Gene Set Enrichment Analysis (GSEA) and Gene Set Variation Analysis (GSVA) were performed separately on GO terms and KEGG pathways to assess functional enrichment associated with this gene’s expression. For GSEA, samples were divided into high-and low-expression groups based on the gene’s median expression value, and enrichment analysis was performed using the clusterProfiler R package^[17]^ to identify enriched KEGG pathways. For GSVA, pathway enrichment scores were calculated for each sample using the GSVA R package^[21]^. Gene sets with p < 0.05 were considered significantly enriched, and results were visualized using enrichment plots and barplots.

### 13. Spatial Transcriptomics Analysis and Dimensionality Reduction

Spatial transcriptomics data from the GSE276841 dataset (generated using the 10x Genomics Visium platform) were processed and analyzed using the Seurat R package (https://satijalab.org/seurat/). Seurat’s standard workflow for dimensionality reduction and clustering. Principal component analysis (PCA) was first applied to reduce the feature space, and Uniform Manifold Approximation and Projection (UMAP) was used to visualize the data in two dimensions.

### 14. Cluster Marker Identification and Cell Type Annotation of Spatial Transcriptomics Data

Spatial transcriptomics data from the GEO database corresponding to a glioblastoma (GBM) tissue sample were processed using Clustering was performed on the PCA-reduced space using Seurat’s default resolution setting, yielding discrete cell clusters. Unsupervised clustering was performed with a resolution parameter of 1.0 to identify distinct spatial clusters. Marker genes for each cluster were then identified by differential expression analysis using a Wilcoxon rank-sum test, applying a log2 fold change threshold of 0.5. These cluster-specific markers were subsequently used to investigate spatial gene expression heterogeneity within GBM tissues. Cell type annotation of these clusters was conducted using the scMayoMap R package^[22]^ with a brain-specific reference dataset. Annotation was performed using default parameters, and the resulting labels were retained without manual adjustment.

### 15. Single-Cell Expression Profiling and CNV Inference

Single-cell RNA-seq data from GBM samples were analyzed using the Seurat R package. Differential expression of the candidate gene was assessed across annotated cell types using Seurat’s FindMarkers function, revealing cell-type-specific expression patterns. Subsequently, large-scale chromosomal copy number variations (CNVs) were inferred at single-cell resolution with the inferCNV tool (default parameters), using non-malignant cell types as a reference population in accordance with standard practice. This integrated analysis enabled a comprehensive evaluation of both transcriptional changes and potential genomic alterations associated with candidate gene.

### 16. KEGG Pathway Activity Inference Using scGSEA and AUCell

We applied scGSEA and AUCell to the GSE276841 spatial transcriptomics dataset from glioblastoma (GBM) to evaluate KEGG pathway activity across individual cells. KEGG gene sets from MSigDB were used for enrichment analysis. scGSEA calculated enrichment scores per cell, identifying significantly enriched pathways (p < 0.05). In parallel, AUCell evaluated pathway activity by computing the area under the recovery curve (AUC) for each cell and labeling high-activity cell clusters based on top-ranked AUC thresholds. Together, these complementary methods revealed distinct spatial patterns of pathway activation at single-cell resolution.

## Results

### 1. Identification of Cellular Senescence-Related Differentially Expressed Genes in Glioblastoma

Seven GBM gene expression datasets were integrated to increase statistical power. After robust normalization and batch effect correction, expression value distributions across all samples became comparable (Figure 2A). The integrated data showed a clear distinction between GBM and normal brain samples, as a heatmap of sample-level expression patterns revealed separate clusters corresponding to tumor and normal tissues (Figure 2B). Differential expression analysis identified hundreds of significantly dysregulated genes (DEGs) in GBM versus normal brain (|log2FC| > 0.585, adj. p < 0.05), including many genes upregulated in GBM and others downregulated, a total of 1246 DEGs are detected (Figure 2C). Overlapping these DEGs with the CellAge database of cellular senescence genes yielded a set of GBM DEGs associated with cellular senescence (Figure 2D). These 129 overlapped senescence-related DEGs represent candidate biomarkers and potential therapeutic targets in GBM.

**Figure 2.**
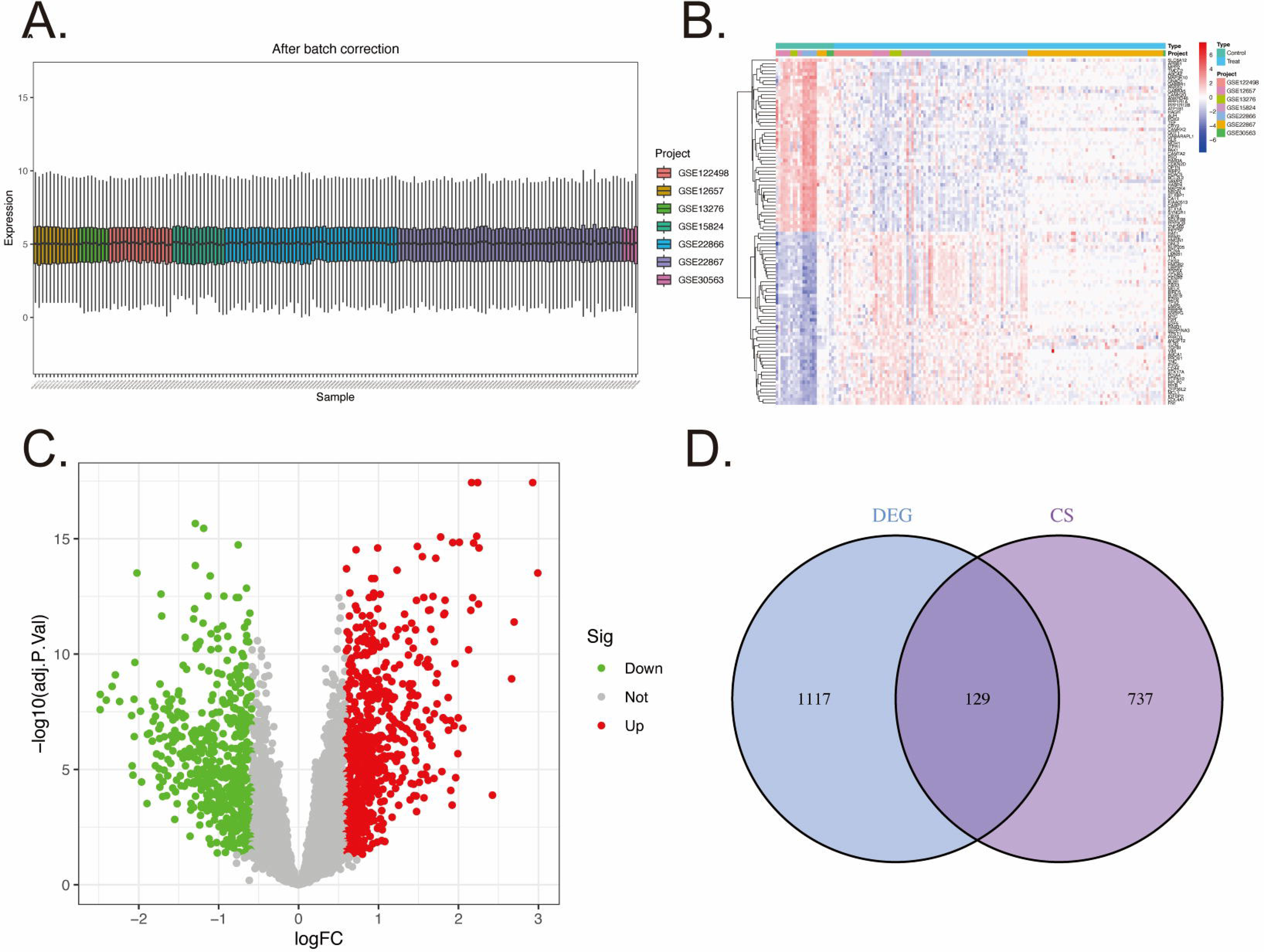
Integrated analysis results with four panels: (A) Barplot of gene expression distributions after normalization and batch effect correction, showing comparable expression distributions across datasets. (B) Heatmap of sample-wise expression profiles after integration, illustrating clear clustering of GBM and normal samples into distinct groups. (C) Volcano plot of DEGs between GBM and normal tissues (significantly upregulated and downregulated genes are highlighted; dashed lines indicate significance and fold-change thresholds). (D) Venn diagram showing the overlap between identified DEGs and CellAge cellular senescence genes, indicating the number of senescence-related DEGs in GBM.

### 2. Functional Characterization of Cellular Senescence-Related DEGs in Glioblastoma via GO and KEGG Analysis

GO enrichment analysis (categorized into BP, CC, and MF) of the CS-related DEGs revealed significant enrichment in cell cycle-and senescence-associated functions. Top BP terms included cell aging, cell cycle arrest, and negative regulation of cell cycle, along with stress responses (e.g., response to oxidative stress) and chromatin organization. Enriched CC terms (e.g., nuclear chromatin, chromosomal region, focal adhesion) suggest many CS gene products localize to nuclear structures or mediate cell adhesion. The top MF terms indicated regulatory or enzymatic roles, including protein kinase regulator activity and transcription corepressor activity. These GO findings imply that CS-related genes govern cell cycle progression, chromatin stability, and stress response programs in GBM (Figure 3A, B). Consistently, KEGG pathway analysis highlighted pathways linked to senescence and tumor suppression. The DEGs were enriched in pathways such as cellular senescence, p53 signaling, cell cycle, and the KEGG “Glioma” pathway, underscoring their relevance to GBM. Other significant pathways (e.g., microRNAs in cancer, transcriptional misregulation in cancer) indicate involvement in broad gene regulatory networks underlying glioblastoma. Together, these enrichment results suggest that the CS-related genes influence critical mechanisms of cell cycle arrest, stress resistance, and GBM progression (Figure 3C, D).

**Figure 3.**
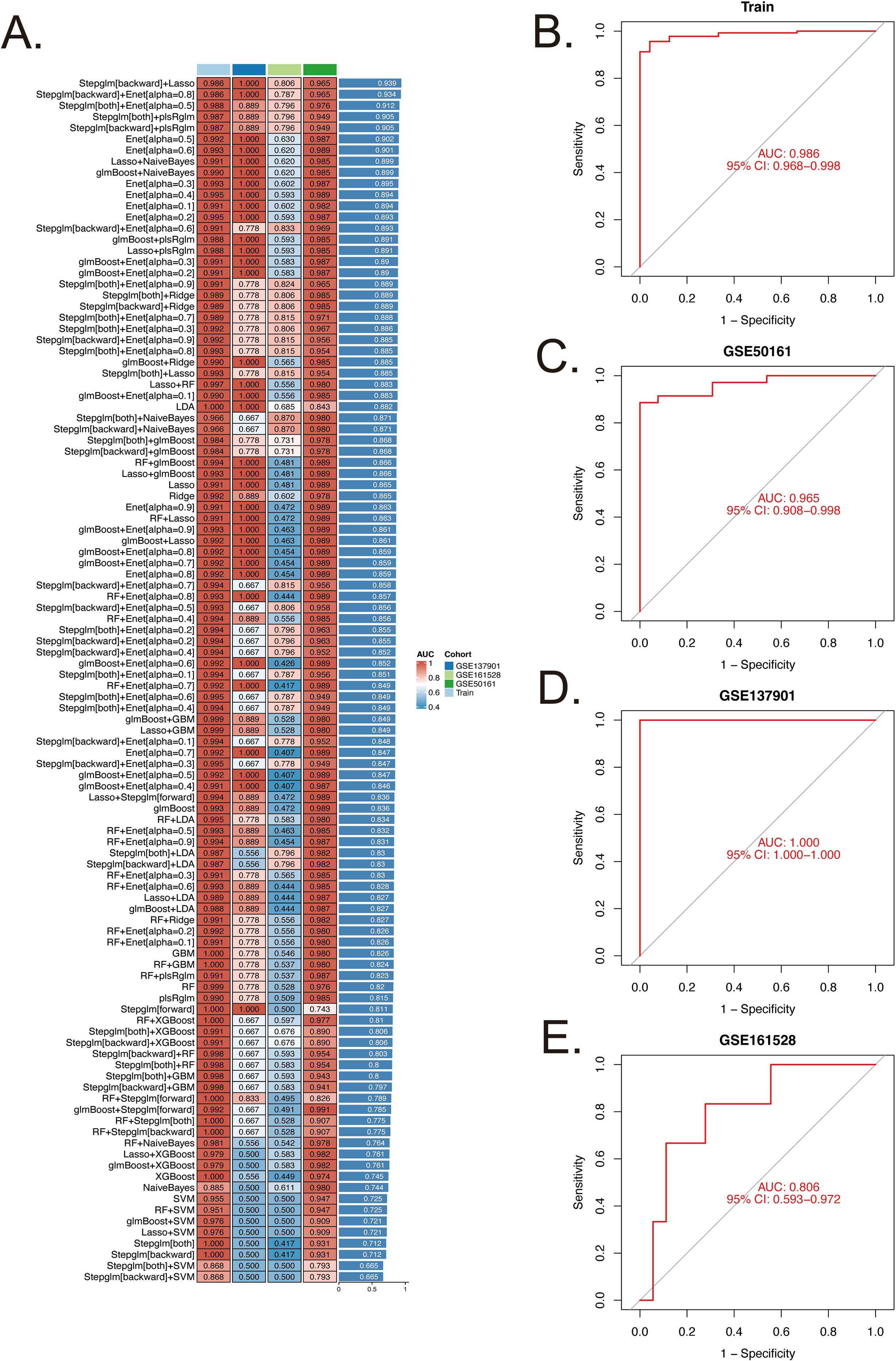
GO and KEGG enrichment analysis of cellular senescence-related DEGs in GBM. (A) GO enrichment bar plot showing the top significantly enriched GO terms across BP, CC, and MF categories; bar length indicates the number of DEGs associated with each term (bars are color-coded by GO category). (B) GO enrichment bubble plot where each bubble represents a GO term (bubble size corresponds to the number of DEGs in that term and color indicates significance level). (C) KEGG pathway enrichment bar plot for the top enriched pathways; bar length reflects the number of DEGs in each pathway. (D) KEGG enrichment bubble plot for pathways (bubble size = number of DEGs; color denotes statistical significance). All displayed GO terms and pathways are significantly enriched (p < 0.05; adjusted p < 1).

### 3. Robust Validation of Integrated Machine Learning-Derived CS-Related Gene Signature in Glioblastoma Cohorts

Among the 113 machine learning models constructed using various single and paired algorithm combinations, the top-performing model (Stepglm[backward]+Lasso) with the highest C-index was selected for further analysis. This model’s CS-related gene signature demonstrated robust and consistent diagnostic value across the training dataset and all three external validation datasets. In the training set, the model achieved an area under the ROC curve (AUC) of 0.92, reflecting excellent discrimination of GBM cases (Figure 4A). Importantly, similar high performance was observed in each independent validation dataset, with AUCs of approximately 0.90, 0.87, and 0.85 in GSE50161, GSE137901, and GSE161528, respectively. The ROC curves for the training and validation sets (Figure 4B–E) illustrate this consistency, indicating that the gene signature maintains high sensitivity and specificity across diverse datasets. These findings highlight the model’s strong diagnostic potential and its generalizability as a biomarker for GBM.

**Figure 4.**
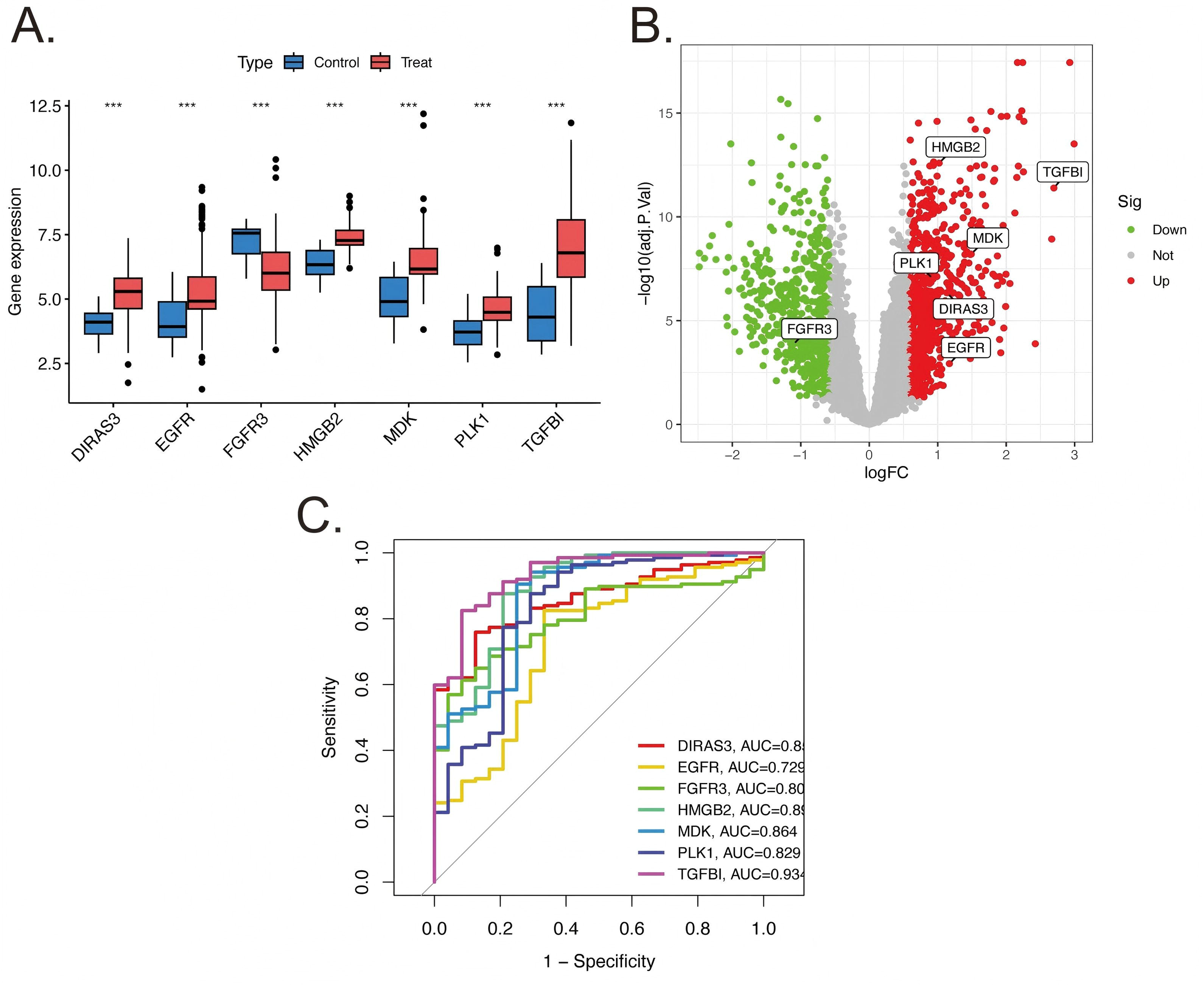
ROC curve analyses of the selected gene signature model in the training and validation datasets. (A) All algorithm AUC value (top-performing model Stepglm[backward]+Lasso, AUC=0.92); (B) Training set (AUC ≈ 0.92); (C) GSE50161 validation set (AUC ≈ 0.90); (D) GSE137901 validation set (AUC ≈ 0.87); (E) GSE161528 validation set (AUC ≈ 0.85).

### 4. Validation and Expression Profiling of Top Candidate Genes From the Optimal Machine Learning Model

Seven Cellular Senescence (CS)-related genes were selected from the top-performing integrated ML model (backward stepwise GLM combined with LASSO) for further validation: DIRAS3, EGFR, FGFR3, HMGB2, MDK, PLK1, and TGFβI. All seven genes were significantly differentially expressed in GBM compared to normal brain (Wilcoxon rank-sum test, p < 0.001 for each). Figure 5A (volcano plot) highlights these candidates, illustrating the magnitude and direction of their dysregulation. Six genes (EGFR, FGFR3, HMGB2, MDK, PLK1, TGFβI) were markedly overexpressed in GBM, whereas DIRAS3 was downregulated relative to normal tissue. Gene-specific boxplots (Figure 5B) further demonstrate these expression patterns, with clear separation between tumor and normal groups. Each selected gene also showed strong diagnostic performance in distinguishing GBM from normal samples. As shown by the ROC curves (Figure 5C), all seven markers achieved high accuracy for classification (each with an area under the curve (AUC) > 0.85). These results underscore their potential as individual diagnostic biomarkers for GBM.

**Figure 5.**
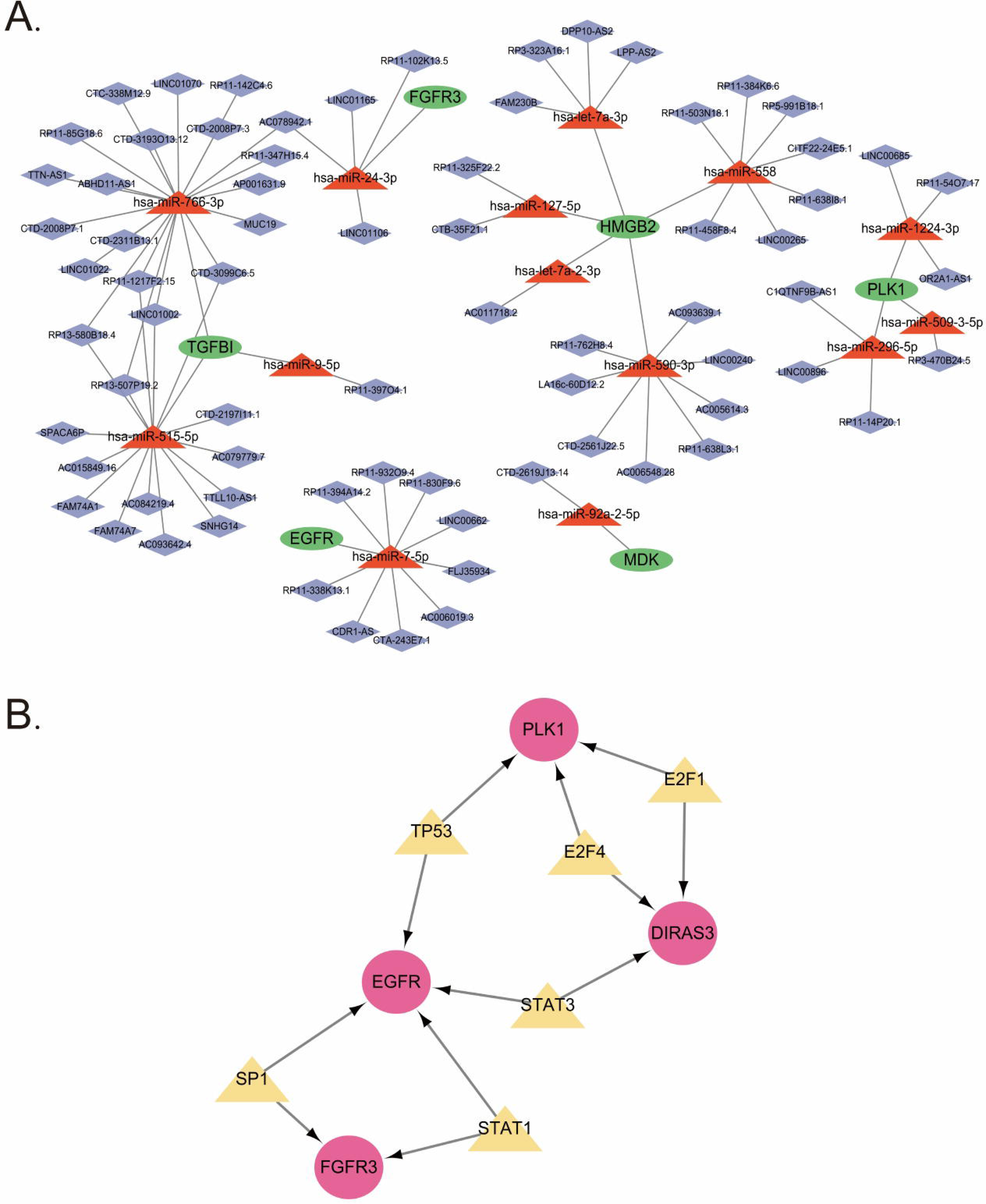
(A) Volcano plot of differentially expressed genes between GBM and normal brain samples. Points to the right of the vertical line (log₂ fold-change > 0) indicate upregulation in GBM, while points to the left (log₂ fold-change < 0) indicate downregulation; all highlighted genes have p < 0.001. (B) Boxplots comparing expression of each selected gene in GBM (tumor) versus normal brain tissue. Each plot shows a significant difference between groups (*** p < 0.001), with higher expression in tumors for EGFR, FGFR3, HMGB2, MDK, PLK1, and TGFβI, and lower expression for DIRAS3. (C) ROC curves evaluating the diagnostic performance of each individual gene.

### 5. GeneMANIA Interaction Network of Candidate CS-Related Genes

Using GeneMANIA, we constructed a gene–gene interaction network for the seven candidate CS-related genes (DIRAS3, EGFR, FGFR3, HMGB2, MDK, PLK1, and TGFβI). The resulting network (Figure 6) shows that these candidates are interconnected through multiple direct and functional associations. GeneMANIA predicted several additional genes in the network as interacting partners or co-expressed nodes, underscoring shared pathways related to cell proliferation and tumor progression. For example, MDK is functionally linked to EGFR signaling, and PLK1 and HMGB2 share associations with cell cycle regulation. The dense connectivity among these genes supports their collective involvement in GBM stemness and progression.

**Figure 6.**
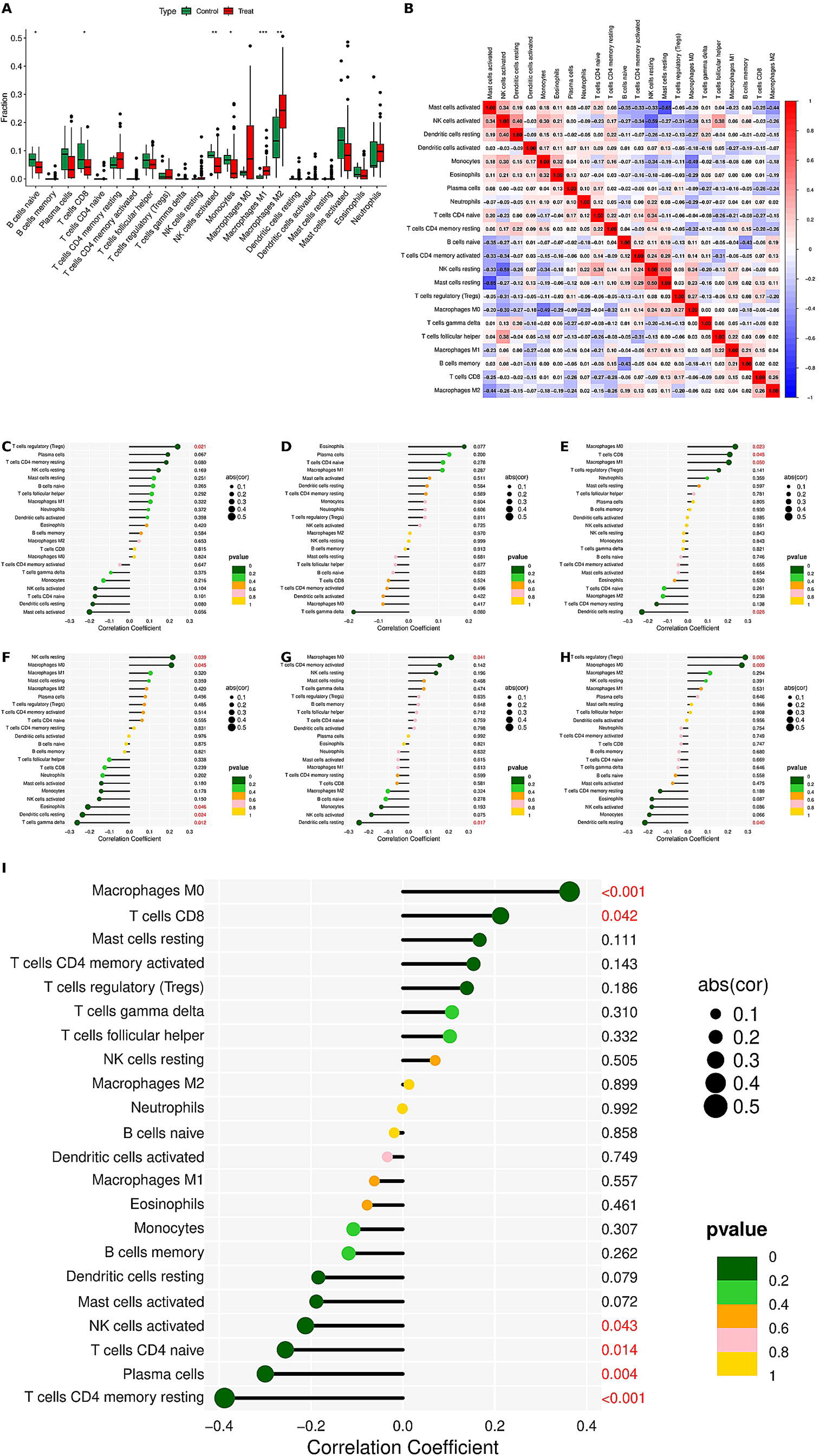
GeneMANIA interaction network of candidate CS-related genes (DIRAS3, EGFR, FGFR3, HMGB2, MDK, PLK1, TGFβI) and predicted interacting genes. Edges represent functional or direct associations.

### 6. CeRNA and Transcription Factor Regulatory Networks of Candidate Genes

A competing endogenous RNA (ceRNA) network was constructed for the seven CS-related candidate genes (DIRAS3, EGFR, FGFR3, HMGB2, MDK, PLK1, and TGFβI) to elucidate their post-transcriptional regulatory interactions. Using starBase predictions (score > 0.8, p < 0.05), we identified multiple lncRNA–miRNA–mRNA interactions for each candidate gene (Figure 7A). Similarly, a transcription factor (TF) regulatory network was assembled based on TRRUST (confidence > 0.7) to map key TFs controlling the candidate genes (Figure 7B). Both networks were visualized with Cytoscape v2.0, revealing complex layers of ceRNA crosstalk and transcriptional regulation centered on the candidate biomarkers in GBM.

**Figure 7.**
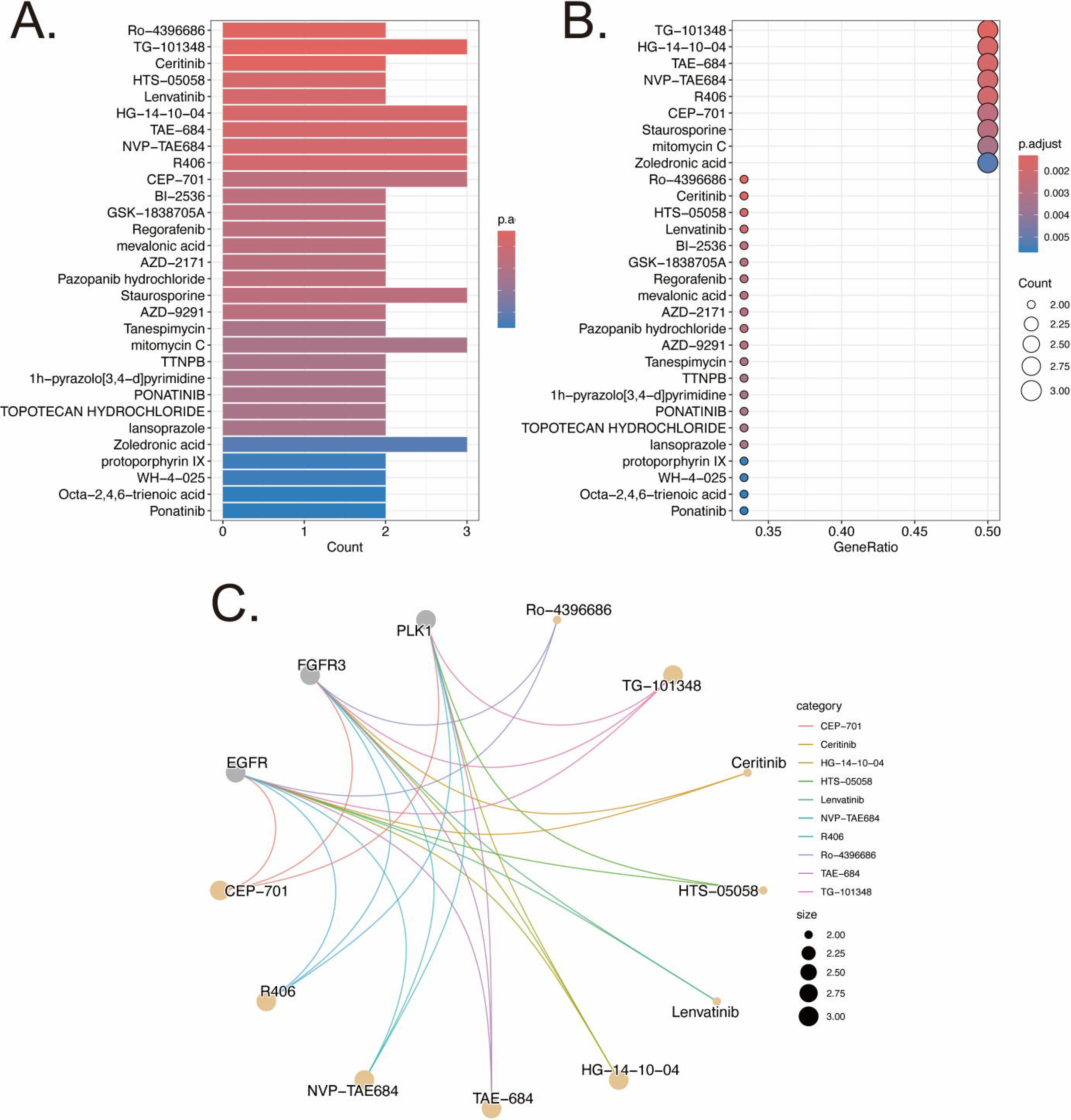
CeRNA and transcription factor regulatory networks of the candidate genes. (A) CeRNA network of the seven candidate genes, illustrating predicted lncRNA–miRNA–mRNA interactions (starBase threshold: score > 0.8, p < 0.05). (B) Transcription factor regulatory network of the candidate genes, showing TF–target interactions from TRRUST (confidence > 0.7). Both networks were visualized using Cytoscape v2.0.

### 7. Immune Cell Profiling and Gene–Immune Interactions in GBM

Using CIBERSORT, we found that the immune cell composition in GBM tumors differs markedly from that of normal brain tissue. GBM samples showed significantly higher infiltration by myeloid-derived cells and lymphocytes associated with immunosuppression. In particular, tumor tissues were enriched in macrophages (especially M2-polarized tumor-associated macrophages) and regulatory T cells (Tregs), whereas normal brain exhibited minimal immune cell presence beyond resident microglia (Figure 8A). A global analysis of immune cell co-abundance revealed that immunosuppressive cell types tend to co-occur: for example, Treg levels positively correlated with M2 macrophage abundance, whereas cytotoxic CD8⁺ T cell levels were inversely related to Tregs (Figure 8B). These shifts underscore an immunosuppressive microenvironment in GBM relative to the largely immune-quiescent normal brain. We next assessed correlations between the expression of seven selected CS-related genes (DIRAS3, EGFR, FGFR3, HMGB2, MDK, PLK1, TGFβI) and immune cell infiltration. Lollipop plots for each gene illustrate distinct association patterns (Figure 8C–I). Notably, higher TGFβI expression strongly correlated with greater macrophage infiltration, whereas DIRAS3 expression showed an inverse relationship with Treg proportions. Several oncogenic genes, including EGFR and MDK, also tended to associate with higher levels of suppressive immune cells, while others had more modest or mixed correlations. The linkET network summary highlights these gene–immune interactions, emphasizing that many of the CS-related genes are embedded in the tumor’s immune landscape (Figure 8J). Together, these results indicate that the identified CS-related biomarkers not only distinguish GBM by their expression but also by their immunological context, potentially contributing to the pro-tumor immune milieu or reflecting immune evasion strategies.

**Figure 8.**
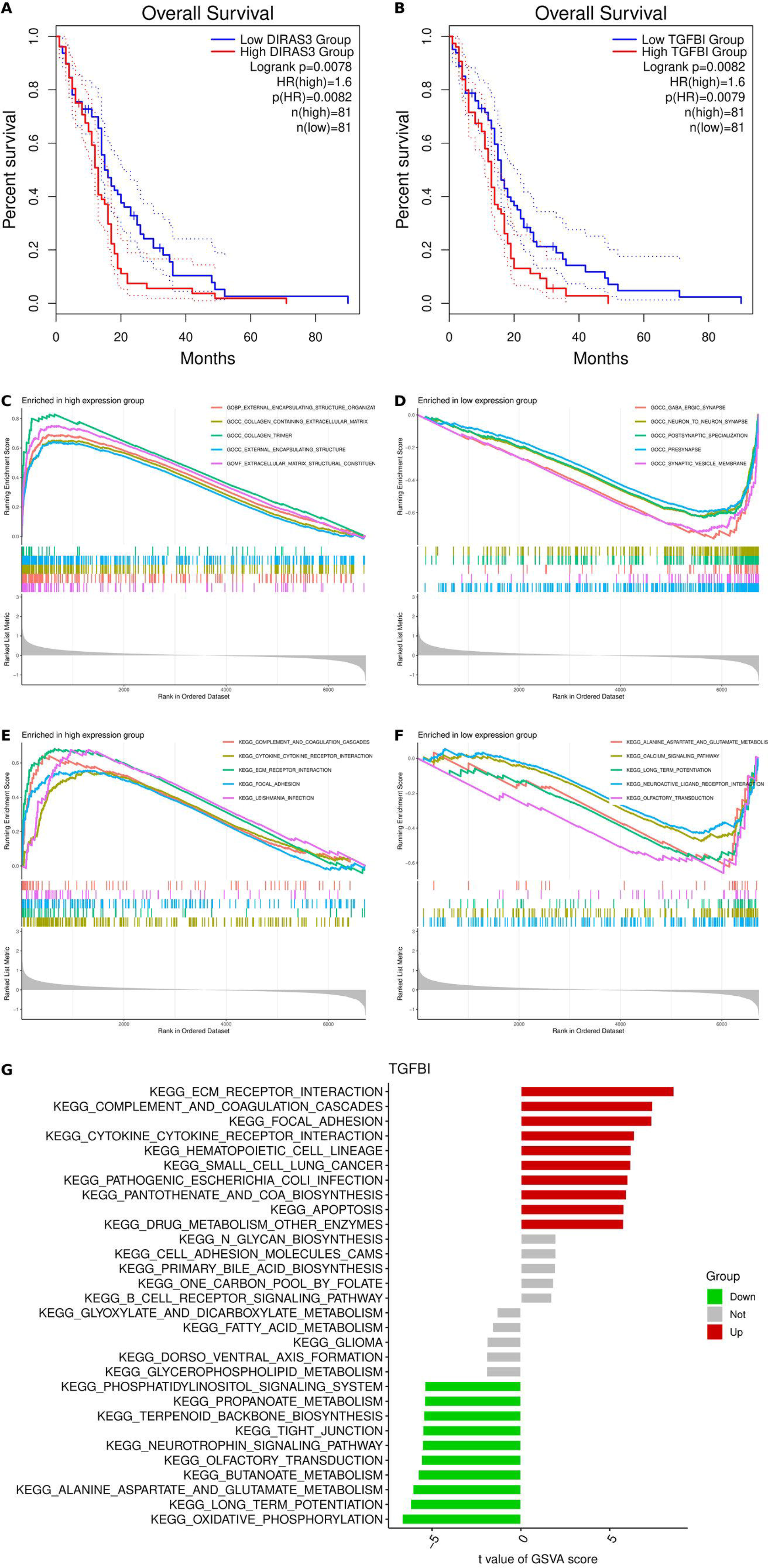
Immune cell profiling and gene–immune correlations in GBM. (A) Relative abundance of 22 immune cell types in GBM tumors vs. normal brain samples (CIBERSORT analysis). Bars indicate mean proportions of each cell type; asterisks denote significant differences between groups (***p < 0.001, **p < 0.01, *p < 0.05). (B) Immune cell co-infiltration heatmap depicting Pearson correlation coefficients among the 22 immune cell fractions in GBM samples (red = positive correlation, blue = negative). (C–I) Lollipop plots showing the correlation between the expression of each CS-related gene and the relative abundance of immune cell types. Each panel corresponds to one gene (C: DIRAS3, D: EGFR, E: FGFR3, F: HMGB2, G: MDK, H: PLK1, I: TGFβI). Lollipop bars represent correlation coefficients with each immune cell type; bar length reflects correlation magnitude and color indicates direction (blue for negative, red for positive correlations). Filled circles denote correlations with p < 0.05. (J) A linkET network diagram summarizing significant gene–immune correlations (p < 0.05). Gene nodes and immune cell nodes are connected by edges, with edge thickness proportional to correlation strength and edge color indicating positive (red) or negative (blue) relationships.

### 8. Drug Enrichment Analysis of Candidate CS-Related Genes

Using clusterProfiler and the DSigDB drug-gene interaction database, we identified multiple compounds significantly associated with the top-ranked cellular senescence (CS)-related genes (p < 0.05, adjusted p < 0.05). As shown in the bar plot (Figure 9A) and bubble plot (Figure 9B), the top enriched candidate compounds include several known anti-cancer and senescence-modulating agents. Each of these top compounds was linked to multiple CS-related gene targets, suggesting a broad impact on the senescence-associated gene network. Notably, the highest-ranking compounds (all FDR < 0.05) target overlapping subsets of the candidate genes. The compound–gene network (Figure 9C) further illustrates that each top drug interacts with one or more of the CS-related genes, highlighting potential multi-target therapeutic avenues for GBM.

**Figure 9.**
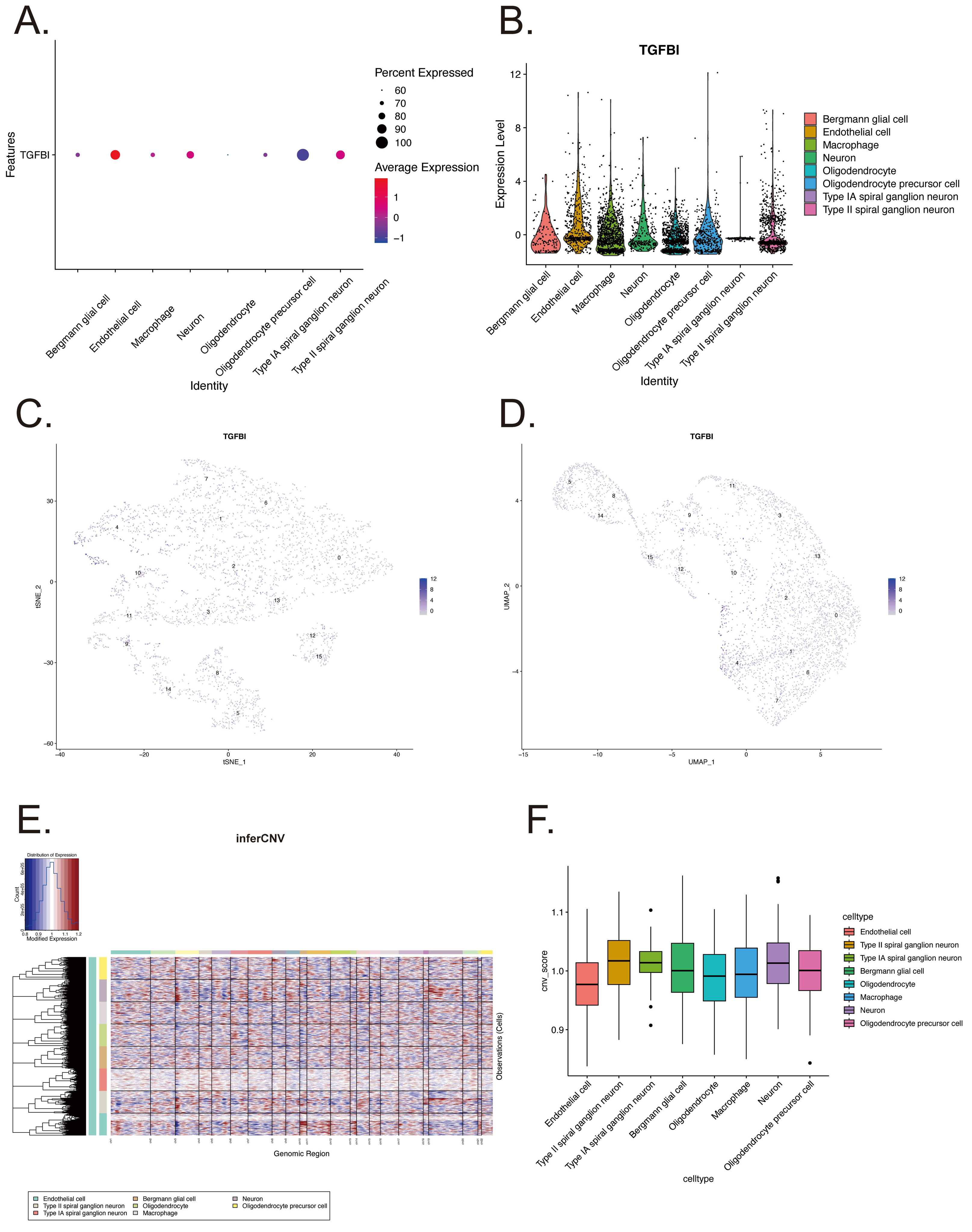
Drug enrichment analysis of candidate CS-related genes. (A) Bar plot displaying significantly enriched compounds (from DSigDB) associated with candidate genes. Bar length reflects enrichment magnitude; all compounds shown meet FDR < 0.05. (B) Bubble plot illustrating enriched compounds. Bubble size represents gene overlap, and color indicates significance (adjusted p-value). (C) Compound–gene interaction network visualizing connections between top enriched compounds and CS-related genes.

### 9. Survival and Functional Enrichment Analysis of TGFβI in GBM

Survival analyses for seven candidate cellular senescence-related genes in GBM using GEPIA2 identified only DIRAS3 and TGFβI as significant (p < 0.05; Figure 10A–B). However, DIRAS3’s Kaplan–Meier curve showed minimal survival separation between high and low expression groups. Therefore, TGFβI was selected for further study. High TGFβI expression was significantly associated with shorter overall survival in GBM (Figure 10B). Gene set enrichment analysis (GSEA) comparing TGFβI-high vs. TGFβI-low tumors revealed significant enrichment of pathways in TGFβI-high tumors. These included Gene Ontology (GO) biological processes related to cell cycle and extracellular matrix organization, as well as KEGG pathways involved in cellular senescence (Figure 10C–F). Similarly, gene set variation analysis (GSVA) confirmed that these pathways were more active in TGFβI-high tumors, which showed greater cell cycle and senescence activity than TGFβI-low tumors (Figure 10G).

**Figure 10.**
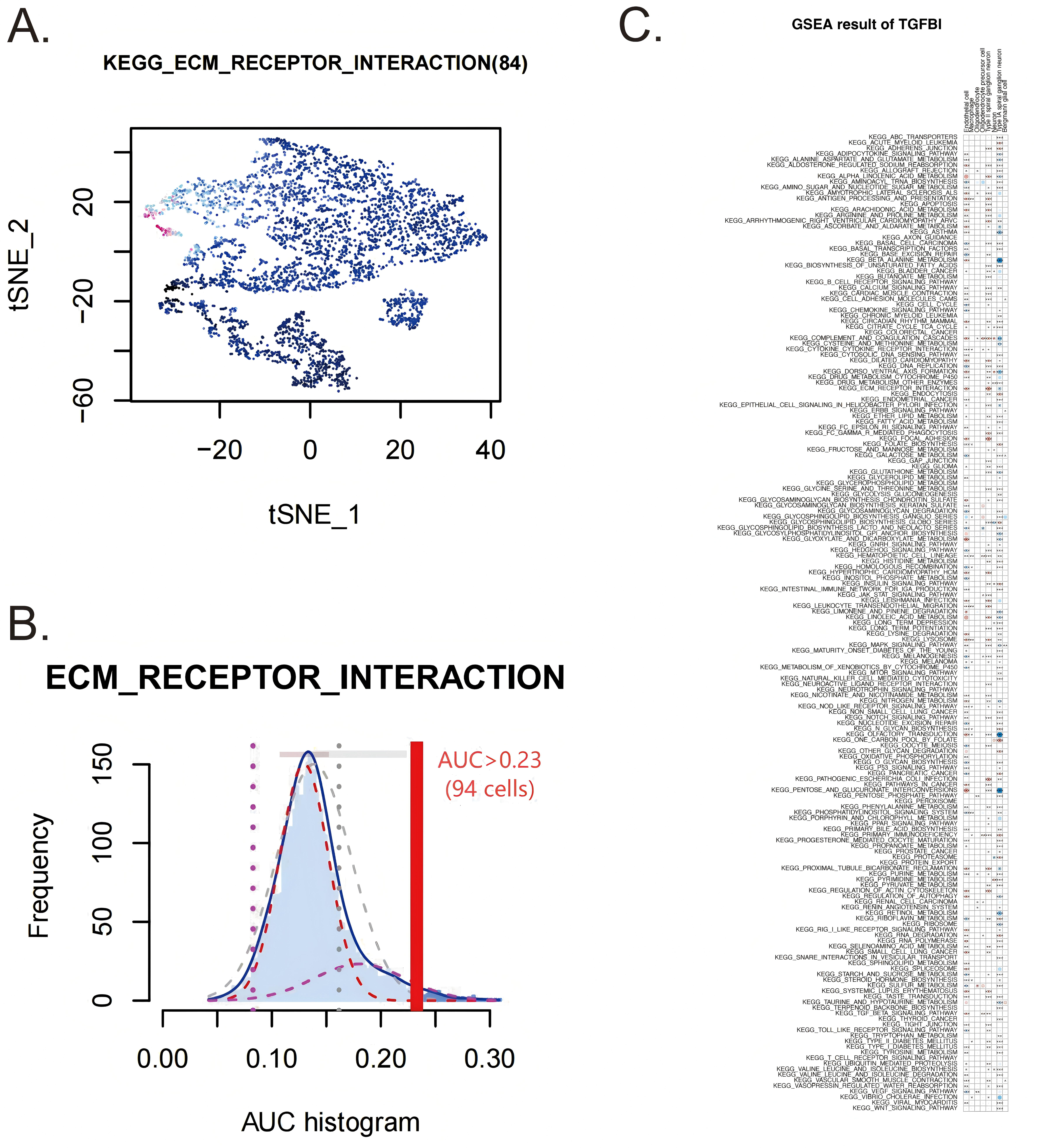
Survival and functional enrichment analysis of TGFβI in GBM. (A) Kaplan–Meier overall survival curve for DIRAS3; although significant (p < 0.05), high vs. low expression groups show minimal separation. (B) Kaplan–Meier OS curve for TGFβI showing significantly shorter survival in the high TGFβI expression group (p < 0.05). (C, D) GSEA highlighting top enriched GO biological processes in TGFβI-high tumors (including extracellular matrix organization). (E, F) GSEA highlighting top enriched KEGG pathways in TGFβI-high tumors. (G) GSVA of pathway activity across GBM samples.

### 10. Dimensionality Reduction and Spatial Clustering of GBM Transcriptomic Data

PCA and UMAP were applied to the GSM8506649 spatial transcriptomic data for dimensionality reduction and visualization. The UMAP projection, colored by unsupervised clusters, revealed multiple distinct groups of spots reflecting diverse cell states (Figure 11A-B). An elbow in the PCA scree plot suggested that the first few principal components captured most of the variance (Supplementary Figure 11A). A heatmap of these top PCs highlights major gene contributors to each component (Supplementary Figure 1B). Mapping the clusters onto the tissue revealed that each cluster occupied a distinct region, indicating spatially segregated transcriptional niches and clear intratumoral heterogeneity. For example, some clusters were enriched at the tissue periphery whereas others localized toward the core. Quality control maps of total UMI counts and detected genes (Supplementary Figures 1C-D) confirmed that the observed spatial patterns were not driven by sequencing depth or gene capture variability.

**Figure 11.**
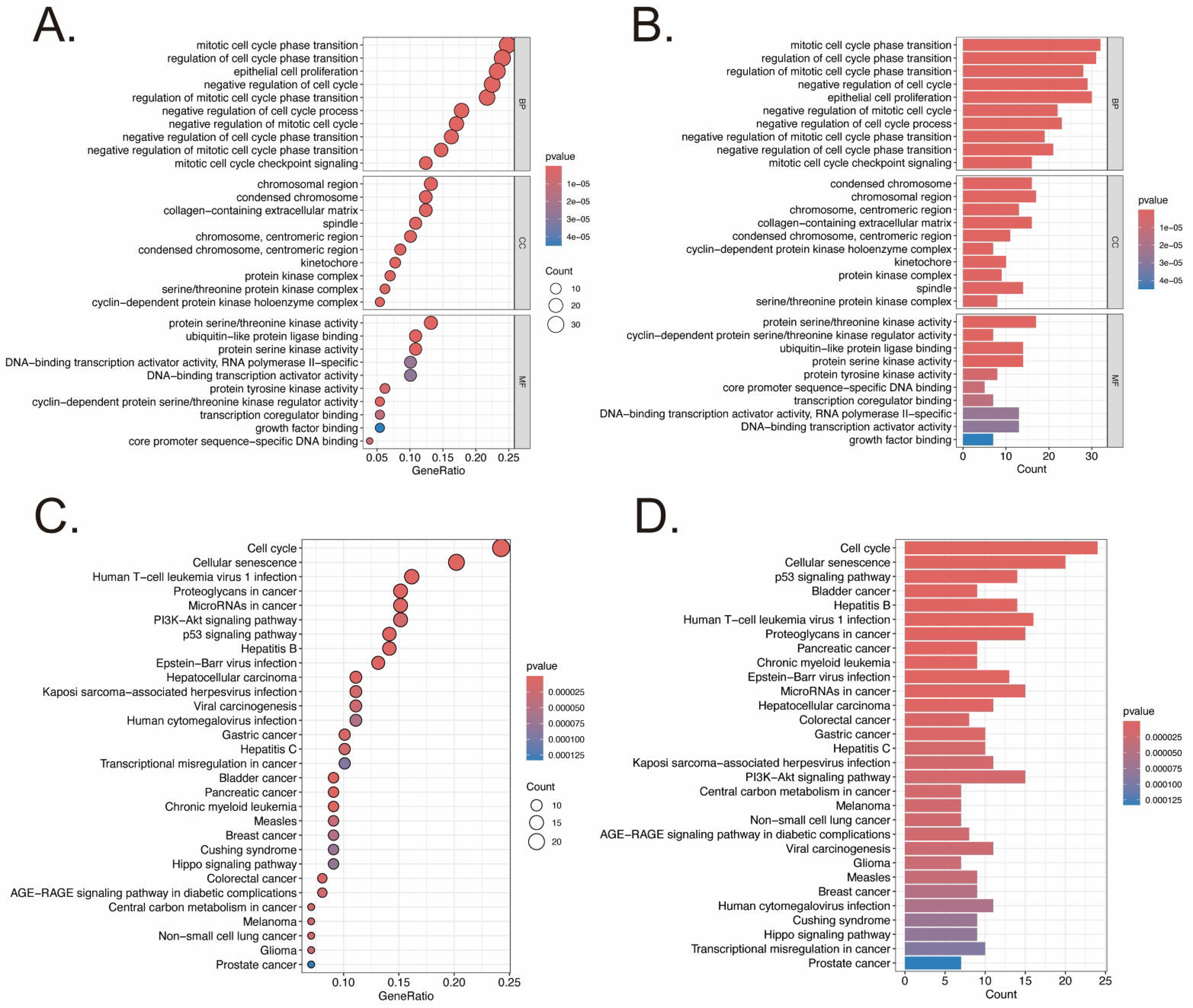
Spatial transcriptomics clustering and quality control metrics. (A) UMAP projection of spots colored by cluster, showing distinct transcriptional profiles. (B) Corresponding tissue section with spots colored by cluster, illustrating spatial distribution of transcriptionally defined regions.

**Figure 12.**
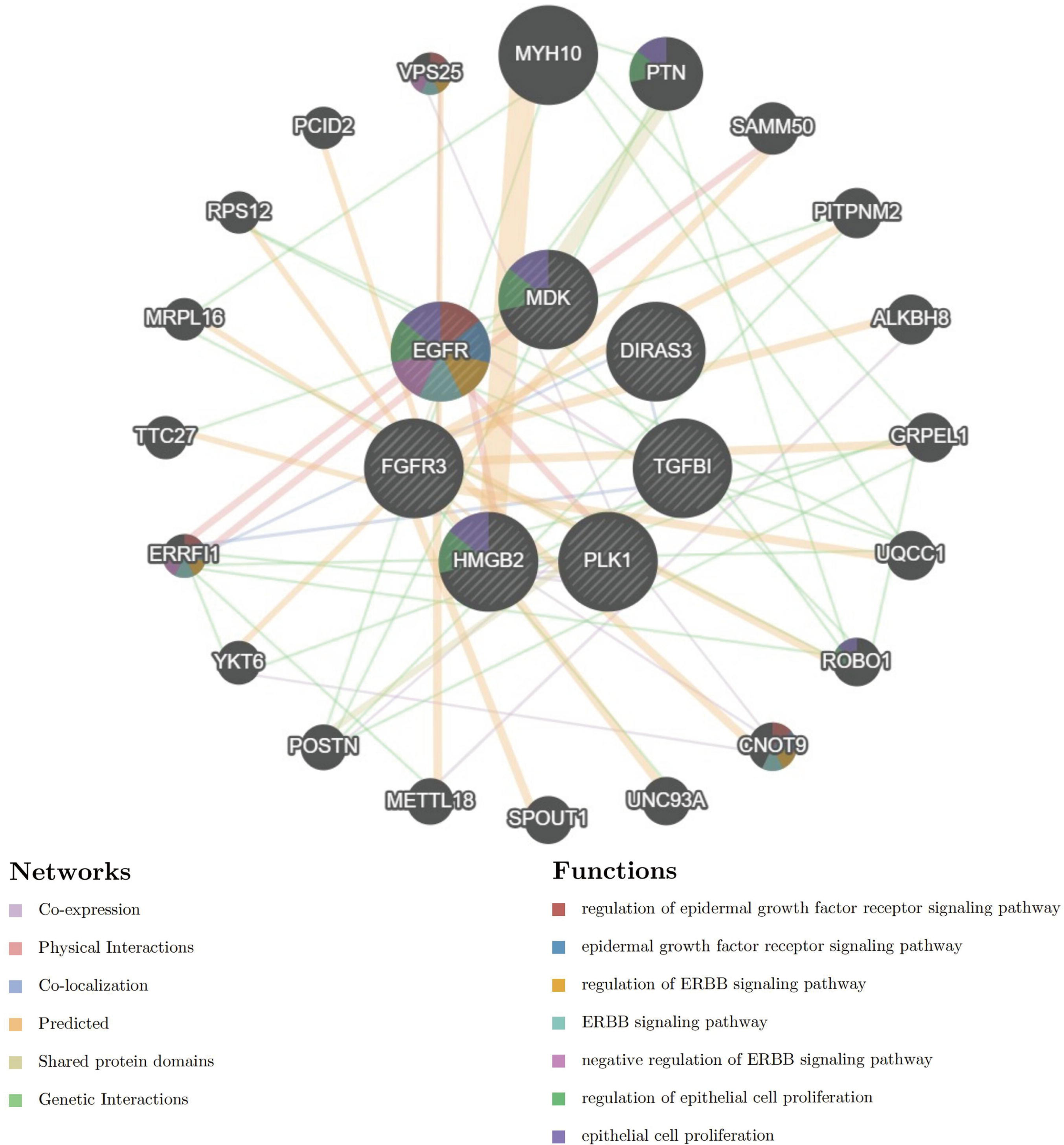
Cell-type annotation of GBM samples using scMayoMap. (A) Dot plot of scMayoMap cell-type annotation scores; dot size and color intensity represent annotation strength, with top annotations highlighted by red circles. (B) Bar plot displaying the proportion of each annotated cell type within GBM sample GSM8506649. (C) t-SNE visualization showing distinct separation of scMayoMap-annotated cell populations. (D) UMAP visualization further illustrating clustering of annotated cell types, highlighting GBM cellular heterogeneity.

### 11. Cell Type Annotation of Spatial Clusters in GBM

Spatial transcriptomics analysis and annotation with scMayoMap identified diverse cellular subpopulations within GBM sample GSM8506649, including endothelial cells, macrophages, oligodendrocytes, astrocytes, and neuronal cells, which localized to specific spatial regions (Figure 11A, B). Dimensionality reduction via t-SNE and UMAP showed clear segregation of these annotated cell types, confirming robust cluster annotation (Figure 11C, D). These findings reveal substantial intratumoral heterogeneity, indicating spatially distinct biological compartments within GBM.

### 12. Single-Cell Expression and CNV Analysis of TGFβI in GBM

TGFβI expression was assessed across all cell populations in the GBM single-cell dataset. A bubble plot summary (Figure 13A) and violin plot (Figure 13B) revealed that TGFβI transcripts were predominantly detected in specific tumor-associated clusters, with minimal expression in non-malignant cell types. Dimensionality-reduced UMAP feature plots (Figures 13C and 13D) further showed that cells expressing high levels of TGFβI co-segregated with the malignant cell population. To investigate whether TGFβI-high cells exhibit genomic aberrations typical of cancer cells, we performed inferCNV analysis using non-malignant cells as a reference. The global CNV heatmap (Figure 13E) demonstrated widespread chromosomal copy number gains and losses in the TGFβI-expressing tumor clusters, whereas reference cell populations displayed flat CNV profiles. Correspondingly, CNV burden scores calculated for each cluster (Figure 13F) were elevated in the TGFβI-high tumor clusters compared to other cells. Together, these results indicate that TGFβI is preferentially expressed in tumor-associated clusters, and that these TGFβI-rich cells co-localize with regions of elevated CNV, suggesting a role for TGFβI in GBM malignant transformation.

**Figure 13.**
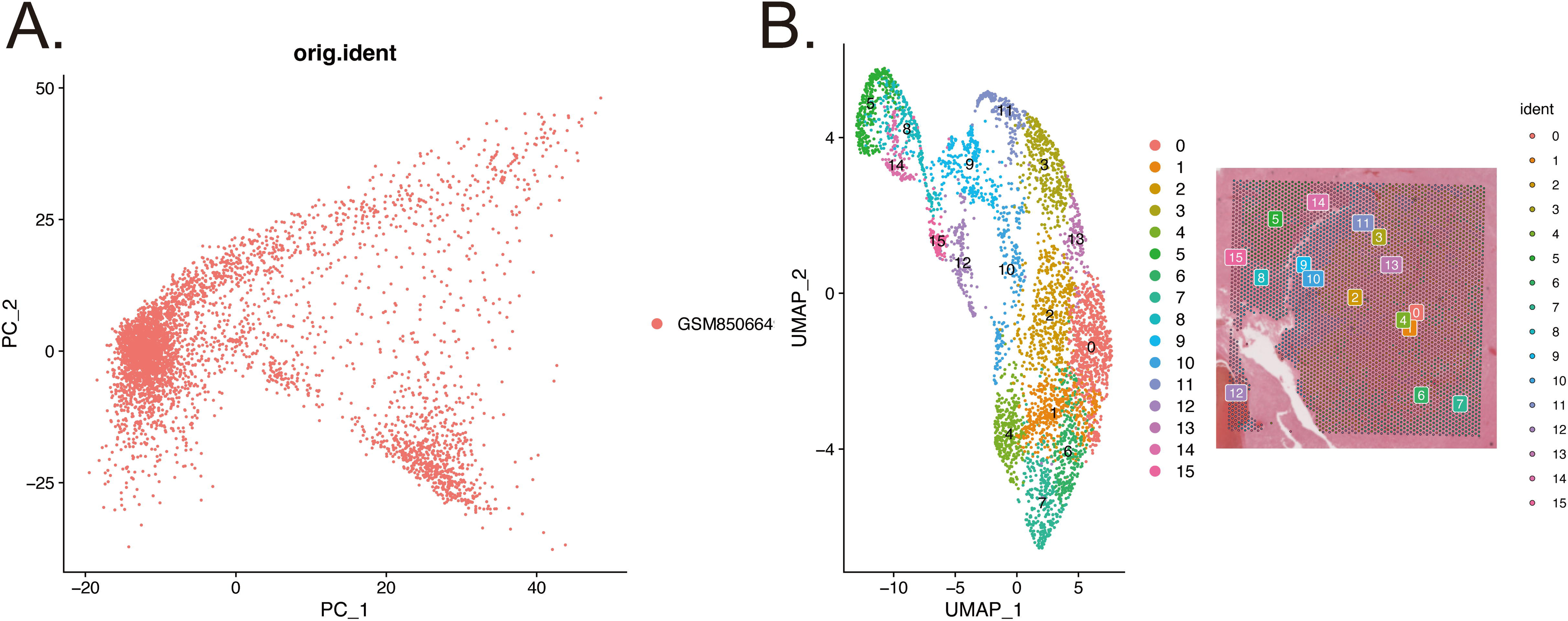
Single-cell expression and CNV analysis of TGFβI in GBM. (A) Bubble plot showing TGFβI expression across all identified cell clusters (bubble size indicates the percentage of cells expressing TGFβI; color indicates average expression level). (B) Violin plot of TGFβI expression levels in each cell type/cluster, highlighting higher expression in tumor-associated clusters compared to non-malignant cells. (C, D) UMAP feature plots of single-cell data with cells colored by TGFβI expression, demonstrating that TGFβI-high cells are localized within the malignant (tumor) clusters. (E) Heatmap of inferred CNVs across single cells (columns) ordered by cell type or cluster, using non-malignant cells as reference. Red and blue color scales indicate relative copy number gains and losses, respectively; extensive CNV alterations are observed in the TGFβI-expressing tumor clusters, whereas non-malignant cells show neutral (baseline) profiles. (F) CNV burden scores for each cell cluster, showing elevated CNV levels in clusters with high TGFβI expression (malignant clusters) relative to non-malignant clusters.

**Figure 15.**
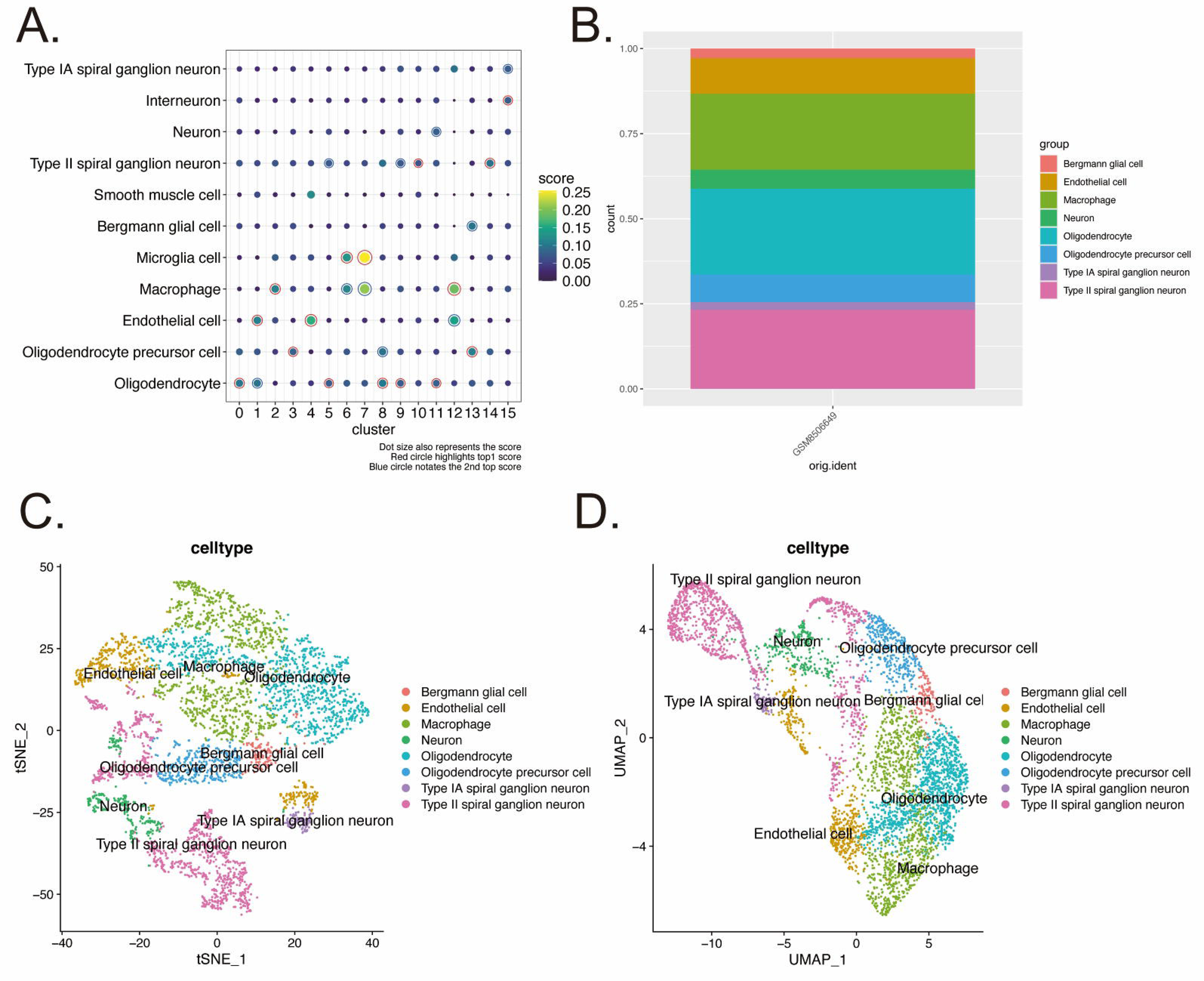
Cellular activity of the ECM receptor interaction pathway in GBM. (A) t-SNE plot of single cells colored by their AUCell AUC score for the KEGG ECM receptor interaction gene set (darker color denotes higher activity). Clusters corresponding to endothelial cells show concentrated high AUC values. (B) Histogram of AUCell AUC scores across all cells, highlighting the score distribution in endothelial cells (blue) versus other cells (gray); a vertical red line indicates the threshold used to define pathway-active cells. (C) Bar chart of scGSEA enrichment scores (NES) for ECM receptor interaction in each cell type. Endothelial cells and type II spiral ganglion neurons show significant positive enrichment (red dot), whereas type IA spiral ganglion neurons exhibit significant negative enrichment (blue dot).

### 13. Single-Cell Pathway Activity Analysis of ECM Receptor Interaction Using scGSEA and AUCell

Using the GSE276841 GBM spatial transcriptomic dataset and KEGG gene sets from MSigDB, we assessed cell-type-specific ECM_RECEPTOR_INTERACTION pathway activity via scGSEA and AUCell. scGSEA identified significant pathway enrichment in endothelial cells, as well as in Type II and Type IA spiral ganglion neurons (Figure 14C). The enrichment scores were positively oriented in endothelial and Type II neurons, indicating upregulation of ECM receptor interactions in these cells, whereas a negative enrichment in Type IA neurons suggested lower pathway activity in that subtype. AUCell likewise highlighted endothelial cells – and to a lesser extent macrophages – as having the highest ECM_RECEPTOR_INTERACTION activity at the single-cell level. A large fraction of endothelial cells exceeded the AUC activity threshold for this pathway, whereas most other cell types showed substantially lower AUC values (Figure 14A and 14B).

## Discussion

The present study harnessed an integrated machine learning framework to unveil cellular senescence (CS)-associated genetic signatures in glioblastoma (GBM). By aggregating 113 models across 11 algorithms, we identified 129 differentially expressed genes (DEGs) linked to senescence (by overlap with the CellAge database) out of an initial 1246 GBM DEGs. This multi-algorithm approach enhanced the robustness of biomarker discovery, mitigating model-specific biases and echoing the growing trend of applying ensemble machine learning to cancer genomics. The identified senescence-related gene set yielded a powerful diagnostic classifier (stepwise GLM + LASSO) with an AUC of 0.92 in training and >0.85 in validation, underscoring the strong signal of these genes in distinguishing GBM. Notably, prior studies have likewise achieved high accuracy by leveraging transcriptomic signatures in brain tumors, reinforcing the validity of our approach^[23]^. These findings position senescence-linked genes as crucial nodes in GBM biology and potential entry points for clinical applications.

Gene enrichment analysis revealed that the CS-related DEGs are enriched in pathways of senescence, cell cycle regulation, and DNA damage response. This is consistent with the biology of therapy-induced senescence in cancer, wherein DNA damage (from radiation or temozolomide) triggers cell-cycle arrest via p53/p21CIP1 and p16INK4a, preventing proliferation. While cellular senescence is classically a tumor-suppressive mechanism, an accumulating body of evidence indicates that senescent tumor cells can paradoxically promote cancer relapse and therapeutic resistance^[24]^. In GBM, residual cells surviving chemoradiotherapy often adopt a senescent-like state instead of undergoing apoptosis. This senescent subpopulation evades immediate cell death and can later “escape” senescence, re-entering the cell cycle to drive tumor regrowth. Clinically, this phenomenon manifests as inevitable recurrence of GBM after standard therapy. Indeed, recent studies have demonstrated that therapy-induced senescent glioma cells contribute to aggressive recurrence, partly by avoiding apoptosis and later regaining proliferative capacity. Our results align with this model: the senescence and DNA damage gene signature likely captures those stress-response programs enabling a subset of GBM cells to endure treatment. Our study’s identification of clear senescence markers in GBM provides a rationale for integrating senolytics into the management of this disease. By removing the senescent, therapy-resistant cell reservoir, one might reduce the likelihood of the tumor’s regrowth.

A striking aspect of the enrichment analysis was the involvement of extracellular matrix (ECM) organization and immune-related pathways among the senescence-associated genes. This suggests that the senescent phenotype in GBM is not cell-autonomous, but deeply connected to the tumor microenvironment. Senescent cells are known to acquire a distinctive secretory profile, the senescence-associated secretory phenotype (SASP), which includes pro-inflammatory cytokines, growth factors, and proteases. The SASP can profoundly remodel the surrounding microenvironment: for example, senescent cells secrete matrix metalloproteinases and other factors that degrade and modify the ECM. This ECM remodeling by senescent cells creates a tissue context that can paradoxically promote tumor invasion and growth. In our data, genes involved in ECM-receptor interaction and matrix structure were enriched, reflecting this pro-remodeling, “wound-healing” signature often associated with senescence. Indeed, senescent fibroblasts in other cancers are known to deposit and reorganize collagen and fibronectin, altering tissue stiffness and signaling to favor malignancy^[25]^. In GBM, such ECM changes at the invasive margin could facilitate tumor cell migration through brain tissue and contribute to the highly infiltrative nature of recurrence.

Our findings dovetail with evidence that therapy-induced senescence in GBM creates a pro-inflammatory microenvironment that attracts tumor-associated macrophages (TAMs) and other immune cells. For instance, after chemoradiation treatment, latent gliomas become enriched in senescent tumor cells and senescent stromal cells, accompanied by increased SASP factors that drive infiltration of macrophages. It has been observed that recurrent GBM tissue (post-therapy) harbors extensive macrophage/microglia infiltration, which is linked to the prior emergence of a senescent, SASP-rich microenvironment. Our study supports this notion: the CS gene signature capturing immune pathways likely represents SASP mediators that promote immunosuppressive myeloid cell recruitment. This is consequential because TAMs are a dominant immune population in GBM that can nurture tumor growth and inhibit anti-tumor immunity. Indeed, TAMs secrete growth factors and proteases (like TGF-β, EGF, MMPs) that support tumor invasion, and they can polarize toward anti-inflammatory phenotypes that dampen T cell responses^[26]^. The SASP of senescent GBM cells likely contributes to such TAM polarization and function. In line with this, we found that macrophage-related signals correlate with our senescence markers, indicating that senescent tumor cells and TAMs form a positive feedback loop in GBM. Senescent cells attract macrophages, and those TAMs in turn can secrete factors (e.g. TGF-β) that reinforce or maintain senescence and mesenchymal transformation in tumor cells. This vicious cycle may underlie the inflammatory but immunosuppressive milieu characteristic of high-grade, therapy-resistant GBM lesions^[27]^.

Among the seven CS-related genes validated in this study, TGFβI (Transforming Growth Factor Beta–Induced protein) emerged as the most prominent biomarker and therapeutic target candidate. TGFβI stood out with a particularly high diagnostic performance (consistent with its individual AUC) and significant prognostic value in GBM patients. This finding is highly compelling in light of accumulating evidence from diverse studies that converge on TGFβI as a crucial mediator of tumor aggressiveness. TGFβI encodes a secreted ECM protein (also known as βIG-H3) that contains an RGD motif and binds to integrins and collagens in the matrix. It is induced by TGF-β signaling, a pathway frequently activated in the GBM microenvironment by both tumor cells and infiltrating macrophages. Elevated TGFβI expression has been correlated with advanced disease and poor outcomes in multiple cancers, and GBM is no exception. A pan-cancer analysis noted that TGFβI is often linked to tumor progression and modulates the tumor immune microenvironment^[28]^. In gliomas, TGFβI expression increases with pathological grade and is highest in the mesenchymal molecular subtype of GBM. Mesenchymal GBMs are known for their invasiveness, inflammation, and therapy resistance, and TGFβI is accordingly a signature gene of this phenotype^[26]^. Consistent with our results, TGFβI is significantly upregulated in glioblastoma samples compared to normal brain and lower-grade tumors, and patients with higher TGFβI levels had markedly shorter survival^[23]^. Thus, our identification of TGFβI as a top biomarker firmly aligns with its known association with aggressive, treatment-refractory GBM.

Mechanistically, TGFβI’s role in GBM appears to be multifaceted, impacting both tumor cells and the microenvironment. One of the most intriguing findings of our study is that TGFβI expression was localized to specific tumor cell clusters characterized by high copy-number variation (CNV) and late pseudotime states, and these TGFβI-high clusters were enriched for ECM-receptor signaling and correlated with macrophage infiltration. This suggests that TGFβI marks a subpopulation of malignant cells, likely the more genomically unstable, advanced tumor cells – that engage in intensive crosstalk with the surrounding stroma. Cells in a late pseudotime state may represent a more differentiated or stressed state of the tumor hierarchy, potentially akin to a mesenchymal or therapy-induced phenotype. Indeed, spatial and single-cell transcriptomic studies have shown that mesenchymal-like GBM cells tend to reside at the tumor margins, assume wound-healing and ECM-remodeling programs, and are accompanied by immunosuppressive myeloid cells^[27]^. Our observation that TGFβI-high cells are associated with macrophages mirrors this pattern. It raises the possibility of a feedback loop wherein TGFβI-expressing tumor cells attract or activate TAMs, and TAM-derived signals (e.g. TGF-β) further induce TGFβI in tumor cells, reinforcing a mesenchymal, senescent niche. In fact, a recent study demonstrated that TAMs can secrete TGFβI in the GBM microenvironment, which then binds to integrin α_vβ_5 on glioma stem-like cells and activates Src/STAT3 signaling to promote tumor growth^[29]^. This TAM-derived TGFβI was shown to enhance the maintenance of GBM stem cells and drive tumor progression, and high levels of TGFβI could be detected in patient serum and cerebrospinal fluid, highlighting its potential as a circulating biomarker. Our findings complement this by indicating that tumor cells themselves (especially in aggressive subclones) also express TGFβI, which likely acts in an autocrine and paracrine fashion to modulate the tumor–TAM interaction. The enrichment of ECM-receptor (integrin) pathways in TGFβI-high cells is in line with the model that TGFβI exerts its pro-tumor effects via integrin-mediated signaling. Notably, integrin–Src–STAT3 signaling is a well-known axis that confers stemness and survival advantages to GBM cells^[30]^. By engaging this pathway, TGFβI effectively connects the senescence/ECM phenotype with core oncogenic signaling in glioma cells.

In summary, our integrated analysis identifies a network of cellular senescence-related genes as pivotal contributors to GBM malignancy, of which TGFβI is a standout player linking therapy-induced tumor cell states to microenvironment remodeling and immune modulation. These findings bridge previously disconnected observations – such as therapy-induced senescence, mesenchymal transition, and macrophage infiltration – into a cohesive model in which senescent tumor cells actively sculpt a pro-tumor niche. Clinically, this advances our understanding of why GBM is so recalcitrant: standard treatments may inadvertently induce a senescent phenotype that tumors exploit for survival and regrowth. Targeting the senescence program, and TGFβI in particular, represents a promising new direction to overcome therapy resistance. As research moves forward, combining senolytic or anti-TGFβI strategies with existing GBM treatments could be the key to disrupting the cycle of recurrence. Our study provides a strong rationale for further preclinical and clinical exploration of TGFβI as both a biomarker and therapeutic target, with the ultimate goal of improving outcomes for patients afflicted with this formidable cancer.

---

Supplementary Figure 1. Quality control and dimensionality reduction of spatial transcriptomics data. (A) PCA scree plot showing variance explained by each principal component (PC). (B) Heatmap of top principal components and their highest-loading genes. (C) Spatial distribution of total UMI counts. (D) Spatial distribution of number of detected genes.

