## Supplementary figures and images for "Screening of Cellular Senescence (CS) Related Genes as Biomarkers and Therapeutic Targets for Glioblastoma (GBM) by Integrated Machine Learning (IML)"

### Supplemental Data 1

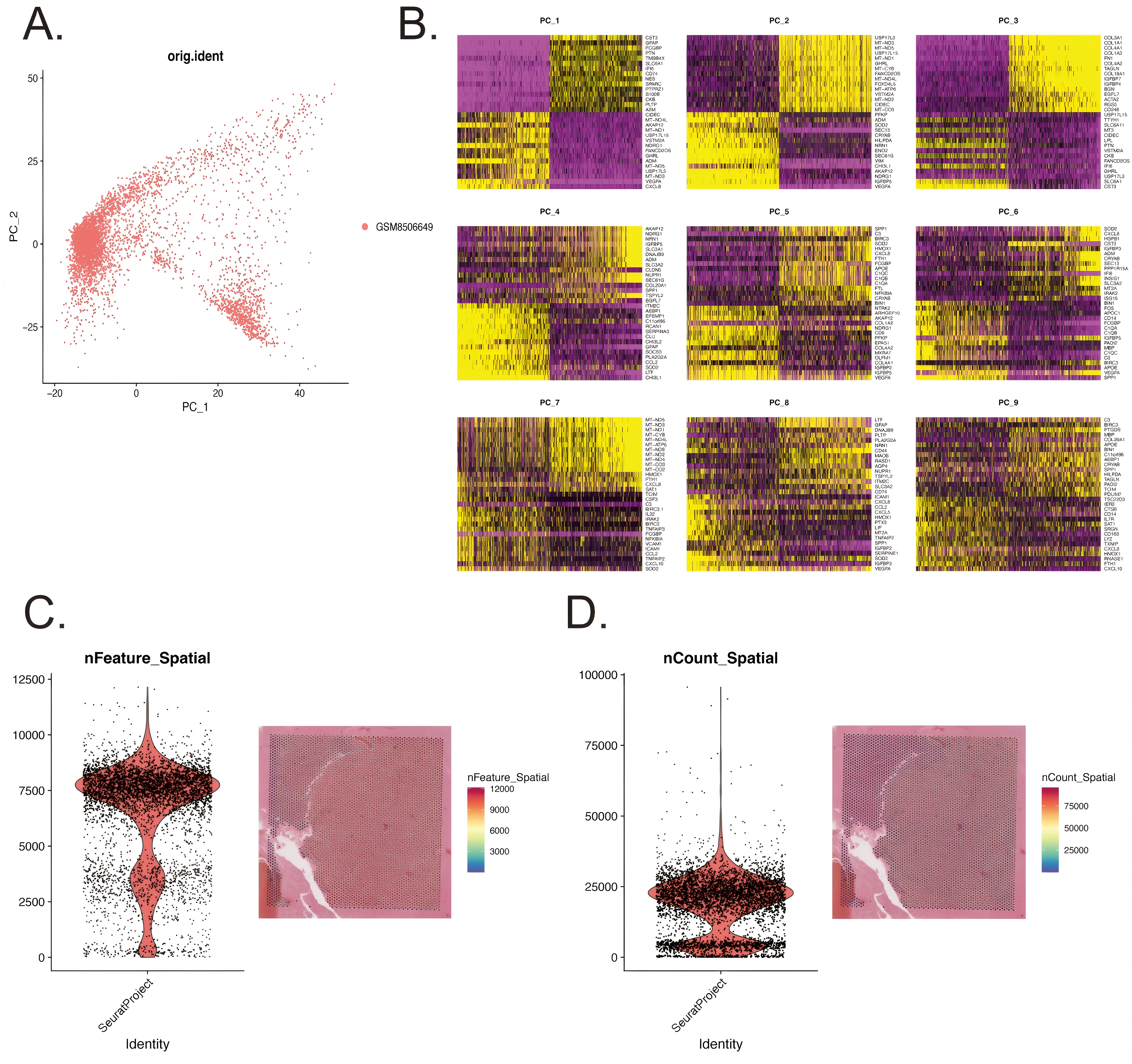
